# Asymmetric oligomerization state and sequence patterning can tune multiphase condensate miscibility

**DOI:** 10.1101/2023.03.11.532188

**Authors:** Ushnish Rana, Ke Xu, Amal Narayanan, Mackenzie T. Walls, Athanassios Z. Panagiotopoulos, José L. Avalos, Clifford P. Brangwynne

## Abstract

Endogenous biomolecular condensates, comprised of a multitude of proteins and RNAs, can organize into multiphasic structures, with compositionally-distinct phases. This multiphasic organization is generally understood to be critical for facilitating their proper biological function. However, the biophysical principles driving multiphase formation are not completely understood. Here, we utilize *in vivo* condensate reconstitution experiments and coarse-grained molecular simulations to investigate how oligomerization and sequence interactions modulate multiphase organization in biomolecular condensates. We demonstrate that increasing the oligomerization state of an intrinsically disordered protein region (IDR) results in enhanced immiscibility and multiphase formation. Interestingly, we found that oligomerization tunes the miscibility of IDRs in an asymmetric manner, with the effect being more pronounced when the IDR exhibiting stronger homotypic IDR interactions is oligomerized. Our findings suggest that oligomerization is a flexible biophysical mechanism which cells can exploit to tune the internal organization of biomolecular condensates and their associated biological functions.

## Introduction

Many cellular reactions are facilitated by colocalization of specific sets of biomolecules into different intracellular organelles such as mitochondria, Golgi apparatus, and endoplasmic reticulum. In addition to the classic membrane-bound organelles (i.e., mitochondria, Golgi apparatus, and endoplasmic reticulum), there are a variety of membrane-less organelles (i.e., stress granule, P-body, and nucleolus) that support diverse cellular functions, from sequestration of translationally stalled mRNA to ribosome biogenesis. Over the last decade, a large number of studies have established biomolecular liquid-liquid phase separation (LLPS) and related phase transitions, as a mechanism for the assembly of these structures, which are typically referred to as biomolecular condensates^1–4^. These findings provide a framework that brings fundamental concepts in thermodynamics and polymer physics to bear on a key aspect of intracellular organization^5–9^.

Past work has shown that the classical Flory-Huggins framework of polymer phase separation is capable of capturing key features of intracellular LLPS^5,10,11 12^. Above the saturation concentration, the enthalpic gain of dispersed biomolecules condensing into droplets outweighs the entropic loss. Flory-Huggins has been helpful for conceptualizing how molecular components of condensates, particularly proteins and nucleic acids with extended and high multivalent interactions capacity, can impact this interplay between enthalpy and entropy to promote phase separation. For example, proteins enriched in condensates often exhibit significant intrinsically disordered protein regions (IDRs), which together with nucleic acid binding partners, have been established as key drivers of condensate formation^13–15^. Such multivalent biomolecular components can be represented as coarse-grained polymers comprised of “sticker” regions interspersed with inert spacers^16,17^. These “stickers” are understood to enable relatively short-ranged favorable interactions and have been recently characterized in purified proteins^10^.

Despite the utility of the Flory-Huggins framework, living cells are far more complex than the inanimate systems it was originally formulated to describe. For instance, cells can dynamically modulate both the enthalpic and entropic factors for regulating the formation condensates, for example through post-translational modifications such as phosphorylation and methylation that tune interactions between sticky regions^18,19^. Living cells also make extensive use of dynamic oligomerization of biomolecular components to modulate LLPS^20–22^. Oligomerization can reduce the entropic cost of LLPS by increasing the effective chain length of proteins, thereby functioning as an entropic knob. Alternatively, monomers can also be potentially sequestered away into stable oligomers, which could hinder LLPS^23^. Oligomerization has also been harnessed to allow the formation of *de novo* condensates^24–27^. In one such system, the optogenetic Corelet technology, light-dependent oligomerization of IDRs and other proteins enabled the quantitative mapping of phase diagrams in living cells, and has been utilized to probe the biophysical properties of condensate nucleation and coarsening behavior^28,29^.

These and other studies demonstrate that although cells are highly complex, fundamental concepts from polymer physics and thermodynamics can be fruitfully employed to understand and engineer intracellular phase behavior. Indeed, dynamic protein oligomerization plays a central role in driving the formation of condensates such as the nucleolus^30,31^, nuclear speckles^32^, and stress granules/P-bodies^22^. Interestingly, these and many other condensates are also often multiphasic, with compositionally distinct phases that are thought to be relevant for their biological functions. For example, the nucleolus is organized into a “core-shell” structure where transcription of ribosomal RNA (rRNA) occurs at the inner “core” (i.e., FC/DFC region), and the nascent rRNA transcripts then undergo sequential maturation steps in the surrounding fluid “shell” (i.e., GC region) before fluxing out of the nucleolus^21,31^. Nuclear speckles also exhibit such a “core-shell” architecture, while stress granules/P-bodies and other condensates provide more complex examples of multiphase organization throughout the cell^33^.

Like most condensates, these archetypal multiphase condensates typically harbor proteins with a significant fraction of IDRs. However, the role of IDRs in driving multiphase organization is not well understood. Recent studies provide indirect evidence that IDRs by themselves might be insufficient to provide specificity for nucleating multiple coexisting phases^22,30,34^, and the molecular mechanisms underlying multiphase formation are poorly understood, particularly within the context of living cells. However, investigating potential mechanisms has proven to be challenging due to the inherent complexity of the intracellular milieu, and a lack of tools for probing phase immiscibility *in vivo*. As a result, how biomolecular components encode their own organization into multiple immiscible subcompartments in living cells remains unclear.

Here, we combine intracellular reconstitution experiments in mammalian and yeast cell systems with coarse-grained molecular simulations to demonstrate that while IDR sequence patterning can by itself dictate multiphase organization, oligomerization greatly amplifies the tendency for segregation into distinct condensed phases. Furthermore, we find that the effect of oligomerization in driving condensate immiscibility is “asymmetric”, with stronger homotypic IDRs showing greater immiscibility when differentially oligomerized. We propose that fine-tuning the oligomerization state of proteins is a mechanism by which cells can modulate the multiphasic organization of biomolecular condensates when the differences in IDR sequence patterning are otherwise insufficiently strong to drive immiscibility.

## Results

### Simulations suggest the role of oligomerization in driving immiscibility

To study condensate miscibility, we first utilized coarse-grained molecular dynamics simulations to examine how oligomerization can drive phase separation. IDRs were modelled as charge-neutral “KE” polyampholytes i.e., polymers comprised of a string of either positively (K)- or negatively (E)-charged beads (Fig. 1A). Building from prior studies^35^, we utilized a set of 30 polyampholytes (SV1-30), representing sequences of increasing charge “blockiness”, quantified by the sequence charge patterning (SCD) metric^36,37^. Consistent with previous studies^34,38^, we find chains that exhibit major differences in their patterning, formed two compositionally distinct phases when mixed (e.g., SV1 and S28, Fig. 1B).

**Fig. 1.**
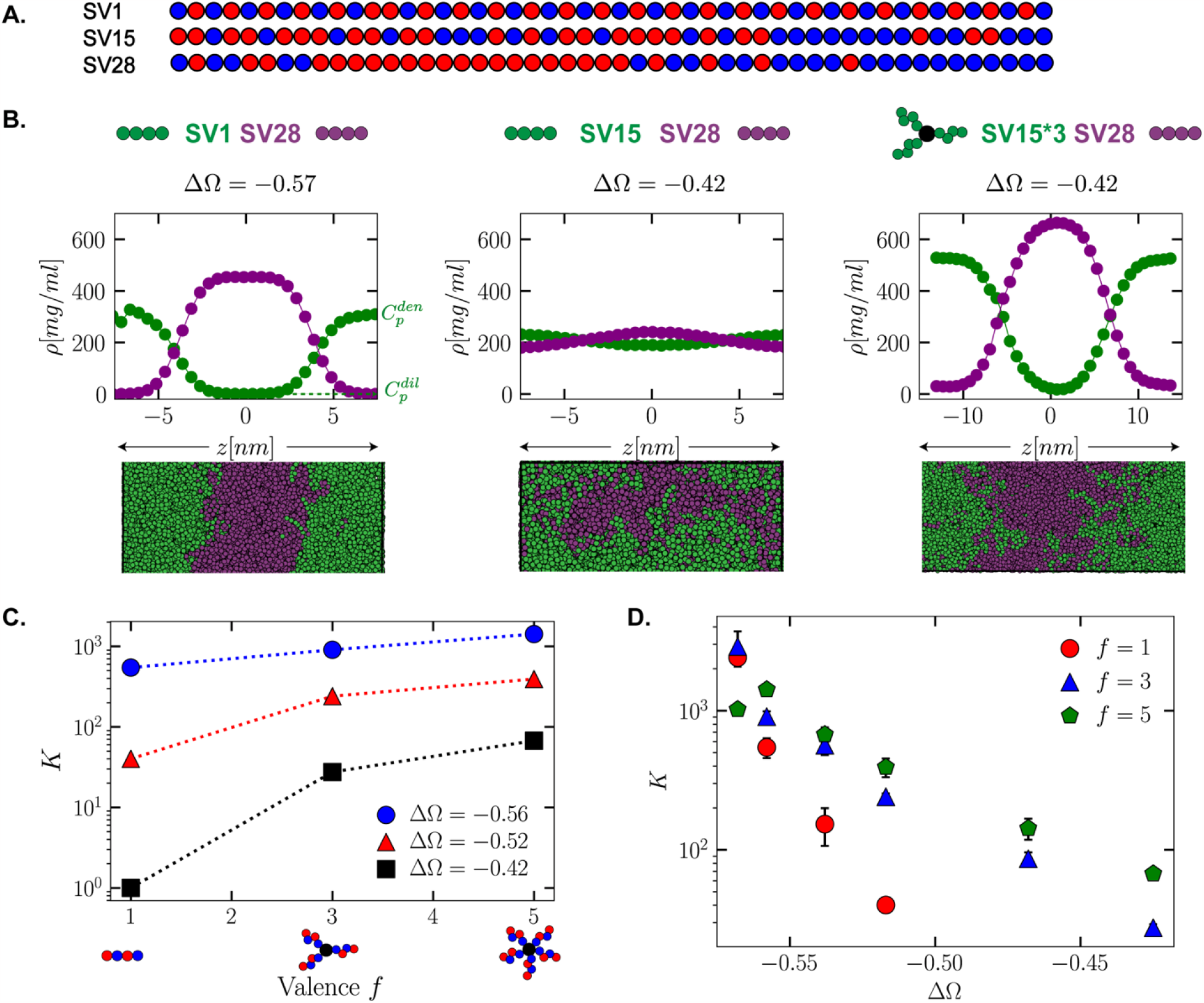
The effect of oligomerization in driving multiphase immiscibility. (A) Model polyampholyte sequences utilized. (B) Simulated density profiles and snapshots of binary mixtures of model polyampholytes highlight the dependence of multiphase immiscibility on the SCD difference (ΔΩ) and oligomerization. (C) Dependence of partitioning on the oligomerization state for 3 different binary polyampholyte pairs. (D) Variation of the partitioning with charge patterning across difference oligomerization states, highlighting how increased oligomerization enhances the range across which polyampholyte pairs can remain immiscible.

To simplify the analysis of the sequence-dependence of this immiscibility, we varied the charge patterning of one of the components *p* while keeping the second component *q* (e.g., SV28) fixed. Furthermore, since two immiscible phases, by definition, will each largely exclude the partitioning of the unfavorable component, we quantify miscibility by examining the partitioning of the component *p*; this mapping is consistent with other immiscibility metrics used in the literature and our own analysis (Supplementary Fig. 1), showing partitioning is a good proxy for phase miscibility. We define the partitioning metric as 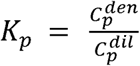, where is the 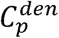 concentration of component *p* in the “*p*-rich” phase while 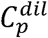 is the concentration in the “*p*-lean” phase. We find the degree of miscibility is dependent on the difference in charge patterning, with larger differences in charge patterning leading to greater immiscibility (e.g., compare Fig. 1B: SV1/SV28 *vs*. SV15/SV28).

In examining how different the sequences must be to drive changes in partitioning/miscibility, we found an unexpectedly strong sequence dependence. For example, reducing the difference in charge patterning -ΔΩ, a normalized form of the SCD metric, by only 36% (Fig. 1B) caused the partitioning to decrease by 2 orders of magnitude. Indeed, our findings suggest that large differences in sequence patterning, and therefore effective interaction strengths, are required for IDRs to form distinct phases. Conversely, IDRs without such differences in charge or sequence patterning might be incapable of demixing into immiscible phases. Thus, the specificity of IDR-IDR interactions alone might, in practice, be insufficient for generating multiple immiscible phases, and instead require additional mechanisms to amplify specificity.

Recent work utilizing synthetic scaffolds has shown that oligomerization can greatly increase the propensity for phase separation by reducing the entropic cost of demixing^2,39^. We hypothesized a similar mechanism might be sufficient for also driving segregation into multiple distinct condensed phases. To examine the role that oligomerization might play in amplifying immiscibility, we simulated binary mixtures where component *p* was modelled as a star polymer with *f* number of arms while component *q* was kept as a single chain. To ensure equal stoichiometry, the mass fraction of the two components was kept equal and two different oligomerization states of f = 3 and 5 were considered. For all cases, we find that increased oligomerization leads to enhanced immiscibility; this can be observed clearly in the case of SV15/SV28 mixtures, which largely intermingle, with little or no distinct phases apparent, while trimerization of SV15 (SV15*3/SV28) gives rise to two distinct phases (Fig. 1B). Immiscibility, as quantified by the fold change in the partitioning, reveals a dependence on both the valence as well as the difference in charge patterning, with more miscible pairs of IDRs having greater fold change in partitioning upon adding valence (Fig. 1C). Simulating a range of binary sequence pairs, we find that higher oligomerization states enable multiphase formation by sequences with relatively similar charge patterning, suggesting that oligomerization effectively amplifies the importance of sequence-encoded interaction preferences (Fig. 1D).

### Light-inducible oligomerization can drive demixing of exogenously expressed nucleolar proteins

To probe the oligomerization-driven demixing hypothesis within living cells, we utilized the light-inducible Corelet system^24,39^ which enables light-triggered dimerization of SspB-iLID and oligomerization of intracellular proteins/IDRs. We co-expressed nucleophosmin-1 (NPM1) fused to SspB (NPM1-mCherry-SspB), together with iLID fused to a multivalent (24-mer ferritin, FTH1) core and a nuclear localization signal (NLS-iLID-GFP-FTH1) (Fig. 2A), in human osteosarcoma (U2OS) cells. Without exogenous oligomerization, the NPM1 construct is strongly partitioned into the nucleolus (Fig. 2Bi). However, upon blue-light activation we observe a striking valence-dependent demixing response. At low overall valence (i.e., ratio of NPM1 concentration to core concentration), we see that the localization of NPM1 remains almost unchanged (Fig. 2Bi). However, for cells with higher valence, the NPM1 exhibits a significant demixing from the nucleolus, and instead becomes enriched at the nucleolar periphery (Fig. 2Bii). Interestingly, at the very highest valence, NPM1 can both demix to the nucleolar periphery, and form new separate condensates in the nucleoplasm (Fig. 2Biii). Taken together, both our simulation and experimental results suggest that oligomerization can cause demixing of condensate components, driving them to form new condensates, or multiphase structures demixed from existing condensates.

**Fig. 2.**
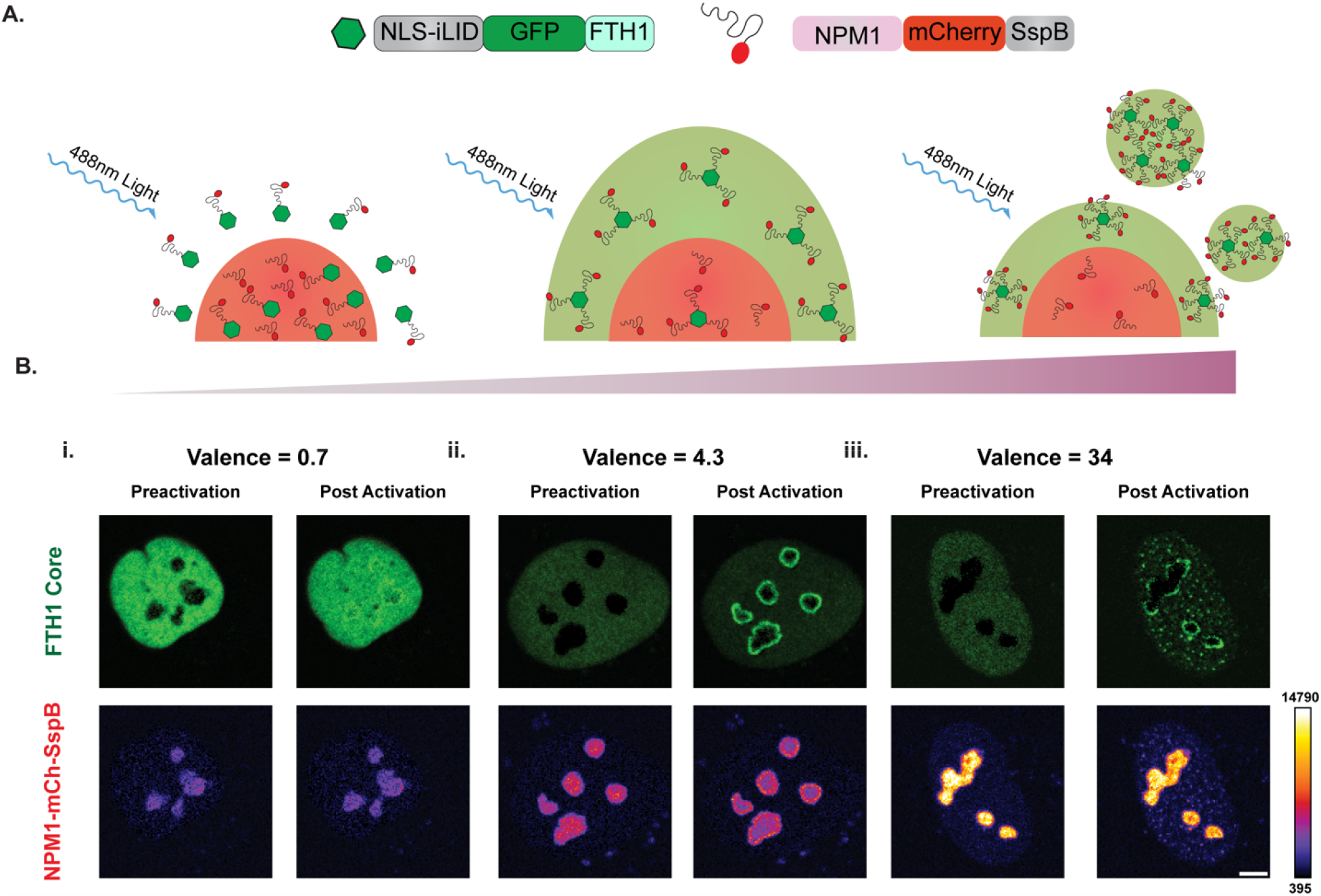
Light induced oligomerization drives demixing of nucleolar localized proteins. (A) Schematics of the optogenetic Corelet experiment in living cells showing oligomerization-driven demixing from the nucleolus. (B) Representative images of the valence dependent changes in nucleolar distribution of NPM1 before and after blue-light activation. The scale bar represents 5 μm.

### Synthetic and orthogonal IDR-core proteins enable the study of condensate immiscibility *in vivo*

The preceding NPM1 data are consistent with our simulations, together showing how condensate components can demix into multiphase condensates upon oligomerization. However, the nucleolus is a highly complex and multicomponent structure, that itself natively exhibits multiphase organization^30^, complicating elucidation of the underlying physics. We thus designed a more tractable intracellular system, which allows for the reconstitution of multiple distinct synthetic condensates in *Saccharomyces cerevisiae*. This approach builds from the Corelet technology (Fig. 2)^39^, but here we utilize a constitutive oligomerization approach, i.e., not light-dependent, so that synthetic biomolecular condensates are always present in cells. Three different cores are used: in addition the previously described ferritin protein comprised of 24 human ferritin heavy chain subunits (FTH1), we also utilize a 60-mer synthetic icosahedron protein I3-01^K129A 40^, and a 24-mer synthetic octahedral protein O3-33^41^. Upon expression in *S. cerevisiae*, mCherry-tagged cores fused to IDRs, such as the N-terminal FUS IDR (FUS_N_) or the C-terminal IDR of heterogeneous nuclear ribonucleoprotein A1 (hnRNPA1_C_), drive the formation of constitutive intracellular condensates (Fig. 3A). This is consistent with previous findings that oligomerization drives the intracellular liquid–liquid phase separation of various IDRs including FUS_N_ and hnRNPA1_c_^39,42,43^

**Fig. 3.**
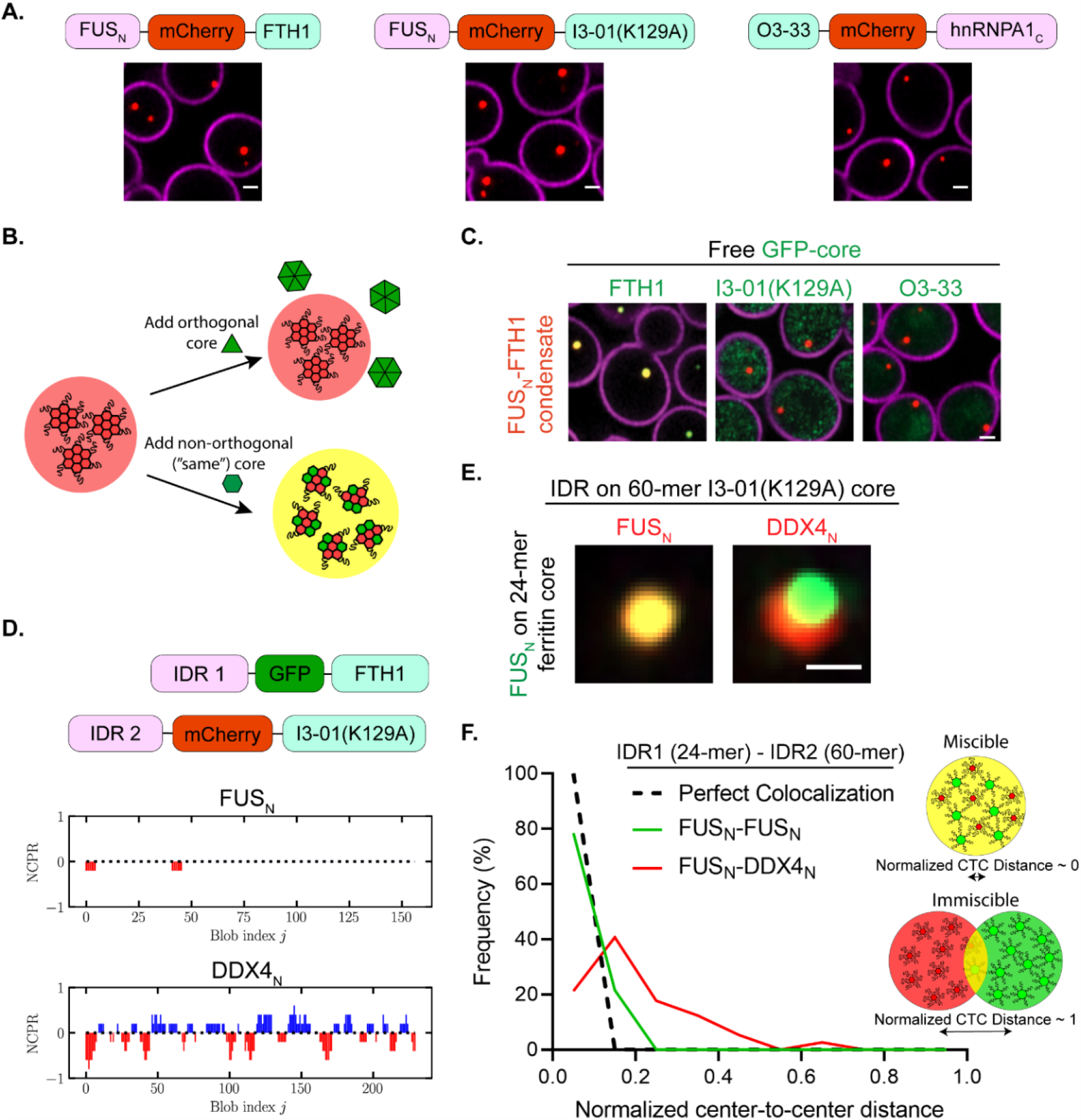
Synthetic and orthogonal IDR-core proteins to study phase immiscibility. (A) Constitutive condensates formed by synthetic IDR-core proteins in yeast. Scale bars represent 1 μm. (B) Schematic diagram of the experiment to examine the orthogonality among different cores (FTH1, I3-01^K129A^, and O3-33). Condensate formed by one core should not recruit other cores if the cores are orthogonal. (C) Only GFP-tagged free ferritin cores are recruited to FUS_N_-FTH1 condensate (I3-01^K129A^ and O3-33 cores are not recruited). The scale bar represents 1 μm. (D) Schematics of the orthogonal IDR-core systems and the residue charge distributions of FUS_N_ and DDX4_N_ are quantified using the net charge per residue (NCPR). The 24-mer FTH1 and the 60-mer I3-01^K129A^ cores are used for studying the interactions between condensate phases formed by different IDRs, for example FUS_N_ and DDX4_N_. (E) Representative images of FUS_N_-FTH1 condensate interacting with FUS_N_/DDX4_N_ -I3-01^K129A^ condensate. The scale bar represents 0.5 μm. (F) The frequency distribution of the normalized center-to-center (CTC) distance between condensates (n > 90). Normalization is obtained by dividing the measured distance between condensates with the sum of radii of the two condensates. The “Perfect Colocalization” is plotted by setting the frequency to 100% for the lowest bin (black dashed line, See Methods for details).

For these different oligomerizing cores to be useful in examining the biophysical determinants of multiphase immiscibility, they must be orthogonal, i.e., each core unit must be capable of self-assembly, independent of the other one. To test for such orthogonality, we expressed GFP-tagged free core subunits (i.e., without any IDRs) in the presence of a different core fused to IDRs; if the cores are orthogonal, then we expect the IDR-core to form a condensate that does not contain the other core. (Fig. 3B). Consistent with orthogonality, we find that IDR-core condensates only recruit subunits of the same core, and when the free core is different from the condensate core, the GFP-tagged core subunits are distributed throughout the yeast cell (Fig. 3C and Supplementary Fig. 2).

We proceeded to test whether these orthogonal IDR-core systems can be used to study condensate immiscibility by using two IDRs (FUS_N_ and DDX4_N_) that have distinct driving forces for phase separation (Fig. 3D). The FUS_N_ prion-like domain contains evenly distributed aromatic residues, that are known to be important for its phase separation^44–46^, while DDX4_N_ phase separation is primarily mediated by electrostatic interactions (cation–pi and pi–pi)^13,15,47^. We find that FUS_N_-FTH1 condensates are not fully miscible with DDX4_N_-I3-01^K129A^ condensates (Fig. 3E), a result that is consistent with recent findings *in vitro*^47^. However, despite their immiscibility, FUS_N_ and DDX4_N_ condensates tend to associate, suggesting some favorable mutual interactions that lead to a reduced surface tension between the two condensate phases. To quantify the degree of miscibility between condensate phases, we measured the center-to-center (CTC) distance between overlapping condensates normalized by the sum of their radii. As expected, the normalized CTC distance profile of the immiscible FUS_N_ and DDX4_N_ pair is markedly different from the profile of a FUS_N_-FUS_N_ control (Fig. 3F). Taken together, our data show that these orthogonal IDR-core systems can be used to study IDR-driven condensate immiscibility, and that significant difference in the sequences of oligomerized IDRs can give rise to condensate immiscibility.

### Oligomerization can drive the miscibility-immiscibility transition of synthetic condensates

In the preceding *in vivo* experiments, we demonstrated that our engineered orthogonal IDR-core proteins can be used for studying the condensate phase immiscibility. To further elucidate the role of oligomerization in tuning condensate immiscibility when the IDRs are not drastically different in their sequence patterning, we sought to use IDRs that share similar driving forces for phase separation. We turned to the C-terminal low-complexity domain of heterogeneous nuclear ribonucleoprotein A1 (hnRNPA1_C_), whose phase separation is driven by aromatic residues similar to FUS_N_ ^10,48^.

We again utilized our orthogonal IDR-core systems in which the 24-mer ferritin and 24-mer O3-33 cores were used to oligomerize FUS_N_ and hnRNPA1_C_, respectively. To obtain varying valence of IDRs, we expressed free ferritin and O3-33 core subunits in the presence of 24-mer FUS_N_-ferritin and 24-mer hnRNPA1_C_-O3-33 condensates (Fig. 4A). Specifically, additional gene copies of ferritin were expressed to reduce the effective valence of FUS_N_, and a set of yeast promoters with well characterized expression strength (strong *TDH3p*, medium *HHF2p*, and weak *RPL18Bp*) was used to reduce the effective valence of hnRNPA1_C_^49^. When the valence of FUS_N_ and hnRNPA1_C_ are both high, we observe two immiscible condensates, a FUS_N_-rich phase, and an hnRNPA1_C_-rich phase (Fig. 4B, C). Interestingly, when the valence of the FUS_N_ condensate is fixed and the valence of the hnRNPA1_C_ condensate is lowered from 24, we observed the two condensate dense phases become miscible (Fig. 4B-D and Supplementary Fig. 3). In contrast, we find that for a fixed hnRNPA1_C_ valence, lowering FUS_N_ condensate valence does not lead to significant change in miscibility (Fig. 4C, D and Supplementary Fig. 3). Taken together, these findings suggest that condensate miscibility can be tuned by the oligomerization state of its protein constituents and, in this case, the oligomerization state of hnRNPA1_C_ is more critical to the miscibility of these two condensates than the oligomerization state of FUS_N_.

**Fig. 4.**
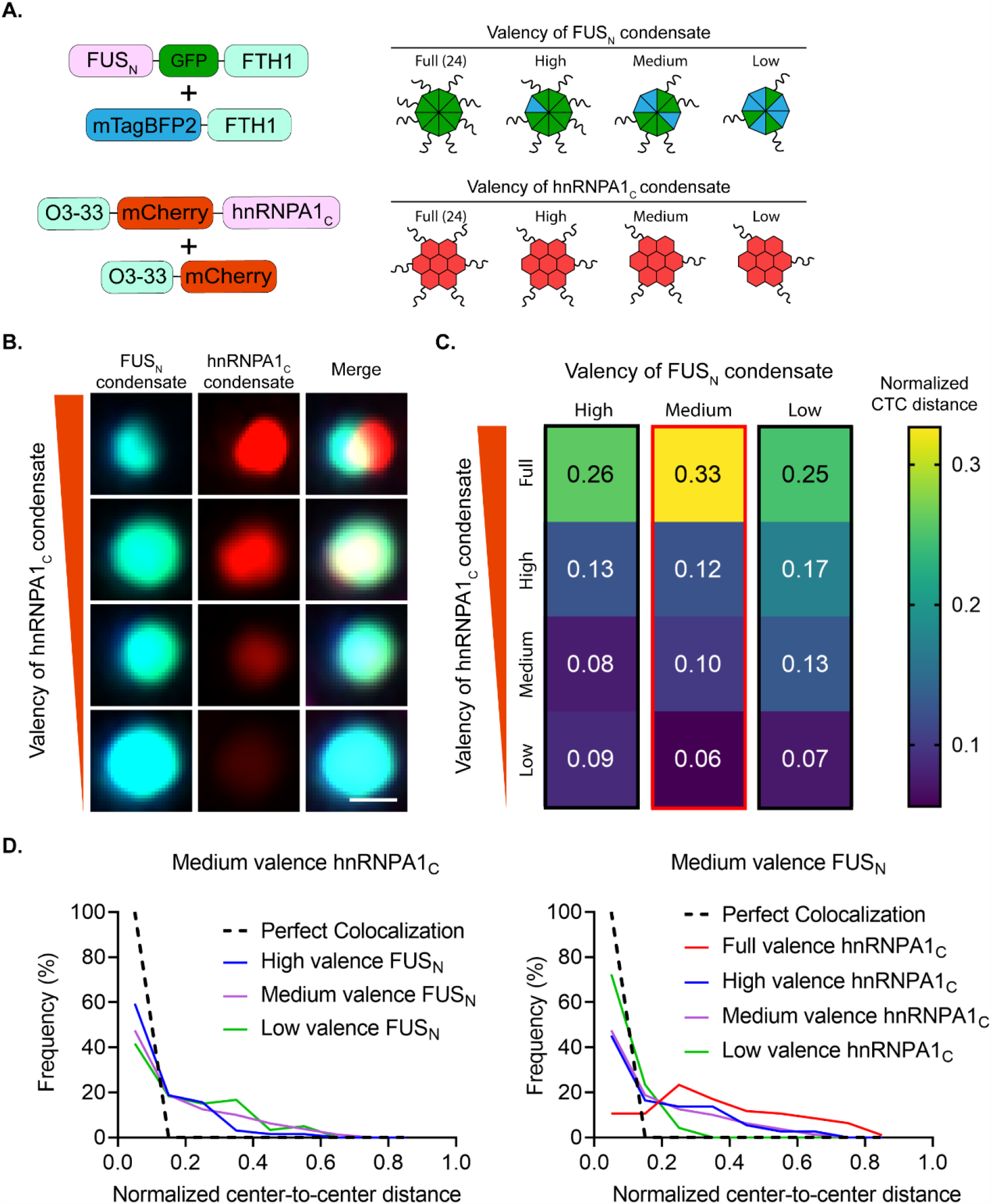
Oligomerization can drive the miscibility-immiscibility transition of synthetic IDR condensates *in vivo*. (A) Schematics of the FUS_N_-FTH1 and hnRNPA1_C_-O3-33 condensates. Both ferritin and O3-33 are 24-mer cores. The valence can be lowered by expressing florescent protein tagged free cores without the IDRs. (B) Examples images of miscibility-immiscibility transition by varying the valence of hnRNPA1_C_ condensate while keeping the valence of FUS_N_ condensate fixed at a medium value (Column with red outline in (C)). The scale bar represents 0.5 μm. (C) The medians of the normalize CTC distance between FUS_N_ and hnRNPA1_C_ condensates at a range of valence (n > 36). (D) Examples of changes in frequency distribution of the normalized center-to-center distance between FUS_N_ and hnRNPA1_C_ condensates when the valence of one IDR is fixed, and the other IDR varies.

### Oligomerization asymmetrically modulates immiscibility between IDR condensate phases

Our finding that the relative oligomerization of hnRNPA1_C_ is more important than that of FUS_N_ to the immiscibility of this system implies some asymmetry in how oligomerization can promote IDRs immiscibility. We find a similar asymmetry in condensate pairs formed by FUS_N_ and DDX4_N._ In particular, we observe clear changes in the CTC distance between the two distinct phases, when one of the two IDRs (FUS_N_ or DDX4_N_) is fused to the 60-mer I3-01^K129A^ and the other to the 24-mer ferritin (Fig. 5A, Bi). To further examine the role of the precise IDR sequence in this oligomerization-driven asymmetric immiscibility, we tested a charge-scrambled version of DDX4_N_ (DDX4_N_CS)^13^ (Supplementary Fig. 4) paired with FUS_N_ or DDX4_N_. One of the most striking examples of this asymmetry is observed with the FUS_N_/DDX4_N_CS pair: when FUS_N_ is highly oligomerized, two distinct phases are observed, while when DDX4_N_CS is more highly oligomerized, no distinct phases are observed. Indeed, condensate phases can either be fully miscible, or immiscible, depending on the relative oligomerization state of the IDRs (Fig. 5A, Bii, Biii). This indicates that whether two different IDR sequences can drive two distinct and immiscible condensates depends both on specific sequence features, and on their *relative* oligomerization states.

**Fig. 5.**
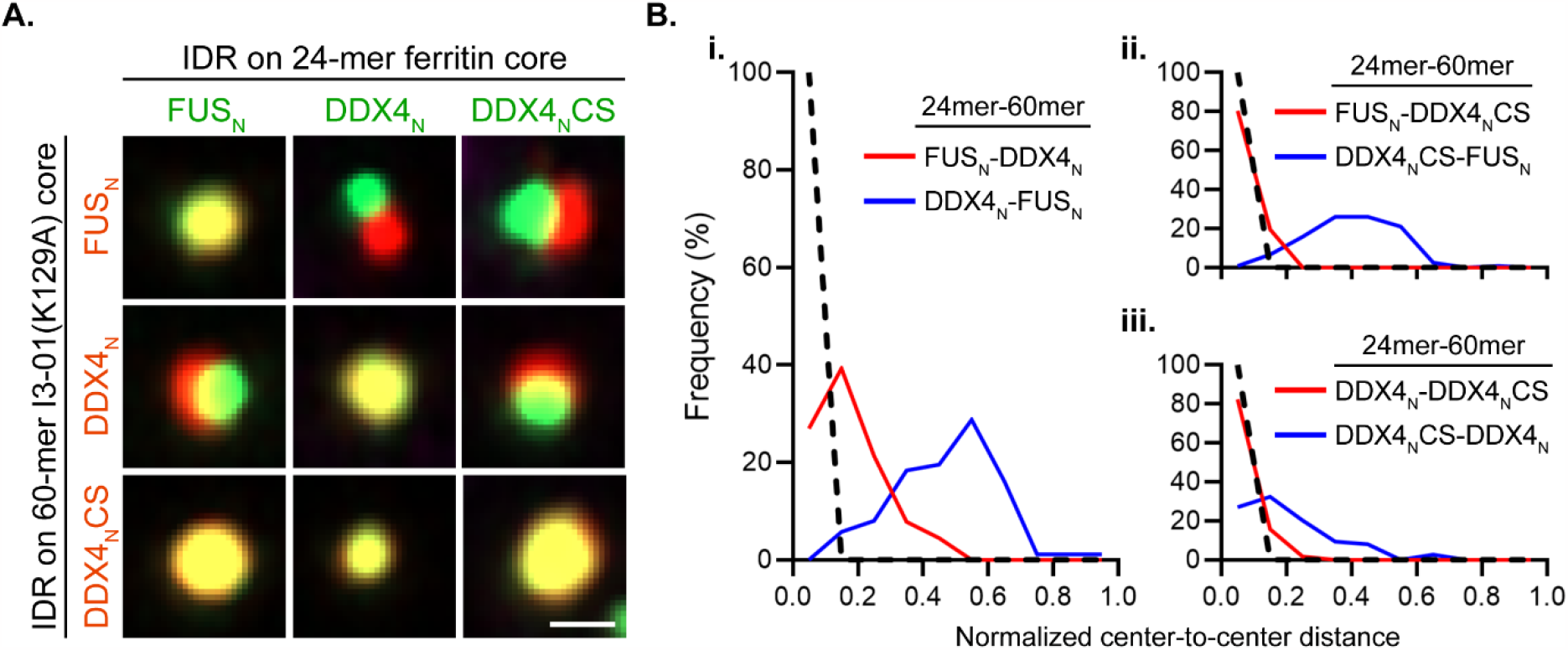
The interplay of oligomerization and IDR sequence patterning in modulating immiscibility *in vivo*. (A) Examples of images of miscible and immiscible condensate phases formed by combination of cores (ferritin and I3-01^K129A^) and IDRs (FUS_N_, DDX4_N_, and DDX4_N_CS). The scale bar represents 0.5 μm. (B) Changes in frequency distribution of the normalized center-to-center (CTC) distance of IDR-IDR pairs when the specific oligomerization state is interchanged (n > 50). CTC distance is normalized by condensate radius (see Methods).

### Asymmetric multiphase formation is a consequence of differential oligomerization states

We noticed that when the miscibility of DDX4_N_/DDX4_N_CS was assayed, a greater degree of immiscibility was observed when DDX4_N_ was more highly oligomerized (a median normalized CTC value of 0.18 *vs*. 0.06) (Fig. 5A, Biii). Since DDX4_N_ is well-established to possess stronger homotypic interactions than DDX4_N_CS^13,15^, taken together this suggests that the entropic “knob” of oligomerization is coupled to an enthalpic driving force. To further examine this physical picture, we returned to our star polyampholyte simulation platform to elucidate how immiscibility depends on the identity of the oligomerized sequence (Fig. 6A). We ran two sets of simulations for each polyampholyte pair, where we studied the effect of oligomerizing each polyampholyte sequence on the miscibility behavior. In the KE polyampholyte model, the strength of the homotypic interaction, as quantified by the critical temperature, has an approximately linear scale with the SCD of the polyampholyte^37^. Thus, polyampholyte sequences with more blocky charge patterns have stronger homotypic interactions. Consistent with our hypothesis, we find that the degree of immiscibility is always higher when the polyampholyte with stronger homotypic interactions (i.e., more blocky charge sequence) was oligomerized (Fig. 6B). For this set of simulations, we utilize the interfacial tension, as a proxy for the degree of immiscibility, which allows us to compare the relative change in miscibility when each IDR in the binary pair is oligomerized. To estimate the strength of the homotypic to heterotypic interaction, we used the ratio of heterotypic to homotypic bonds *<*_*B*_ formed by the star polyampholytes as a proxy. In confirmation with our interfacial tension calculations, we find that the bond ratio (*<*_*B*_) was consistently higher when the more homotypic IDR in the pair was oligomerized. These results suggest that oligomerization enhances the effective strength of homotypic interactions and penalizes the formation of heterotypic bonds. This energetic cost is significantly higher when the oligomerized polyampholyte has strong homotypic interactions, thereby leading to an asymmetric effect.

**Fig. 6.**
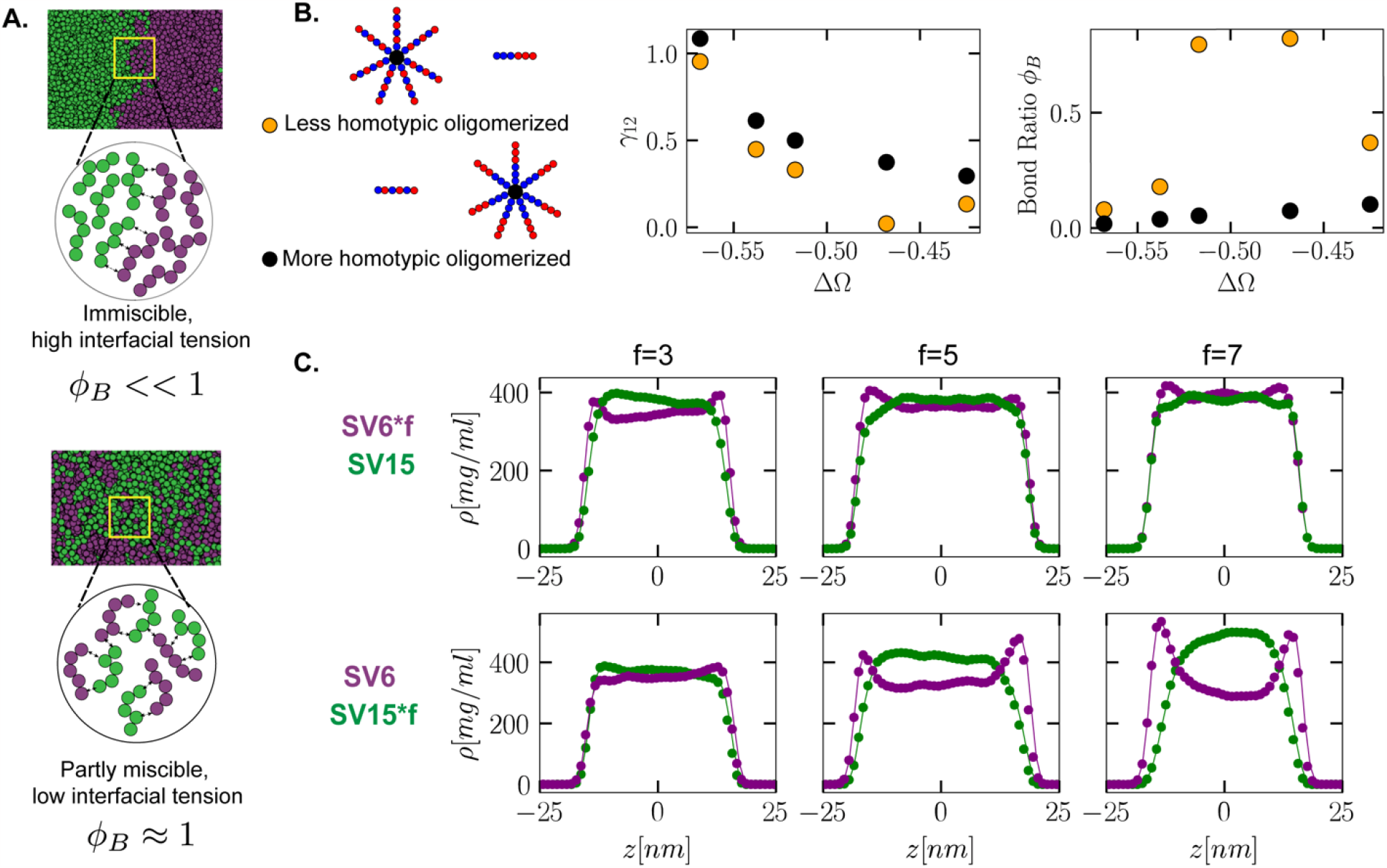
Asymmetric condensate immiscibility can arise from differential oligomerization. (A) Snapshots of strongly and weakly immiscible two-phase systems demonstrate the relative variation in their interfacial tension. Schematics show how fewer heterotypic bonds are formed for the strongly immiscible system. (B) The interfacial tension, utilized as a measure of immiscibility, is consistently higher when the more homotypic IDR in the pair is oligomerized. The bond ratio defined as ratio of unlike to like noncovalent bonds formed by the oligomerized IDR which is always lower for the system with the more homotypic IDR oligomerized. (C) The binary mixture of SV6 and SV15 demonstrates how oligomerization asymmetrically tunes multiphase formation.

Based on our findings, we reasoned that even polyampholytes with small charge patterning differences should be asymmetrically immiscible at sufficiently high valences. To test this hypothesis, we simulated the binary pair SV6 and SV15. We find that when SV6, which has weaker homotypic interactions, is oligomerized, the two components are perfectly miscible (Fig. 6C, top row). In contrast, when SV15 is oligomerized, the two component phases begin to demix especially at higher valences (Fig. 6C, bottom row). Thus, while oligomerization can serve to amplify small sequence-dependent differences in interaction preference that underlie phase immiscibility, the impact of this effect depends on which sequence is oligomerized.

## Discussions

The molecular mechanisms behind the formation of biomolecular condensates and their resultant morphology and material properties have been the subject of intense study over the past decade. Most of this work has been directed toward understanding how biomolecular components in the cellular milieu can condense into single liquid-like phases. Yet, some of the most well-established endogenous condensates (e.g., nucleolus) are organized into multiple immiscible coexisting phases. Recent work has established that the organization of these coexisting immiscible phases is controlled by differences in interfacial tension, a mesoscale property, between the different phases^30^. However, the molecular mechanism underlying such mesoscale organization remains poorly understood, especially *in vivo*. In this study, we sought to investigate the role of oligomerization as a potential general molecular mechanism for driving the formation of multiphasic condensates.

To probe the biomolecular driving forces behind multiphase formation, we engineered orthogonal scaffolding “cores” of distinct valences (24-mer and 60-mer) that enable the formation of synthetic condensates *in vivo*. The different valence state of the cores allowed us to probe the influence of oligomerization on the miscibility between distinct condensates, which had not been studied previously. By fusing different archetypal IDRs to the cores, we found that the propensity to form multiple immiscible phases depends on the balance of sequence-encoded interactions. These findings suggest biomolecular condensate components feature evolutionarily tuned sequence determinants that localize proteins to specific subcompartments.

A key unexpected finding, clearly seen in both our experimental and computational investigations, is that multiphasic structure depends asymmetrically on the oligomerization state of IDRs. For all IDR pairs tested, a greater degree of immiscibility occurs in configurations in which the more homotypic IDR is fused to the higher valence core. Our simulations also revealed that oligomerization increases the propensity to form multiphase structures, by amplifying the relative difference in the strength of homotypic *vs*. heterotypic interactions. This leads to an emergent asymmetry in which IDR is oligomerized, with a stronger driving force for multiphase organization when more homotypic IDRs are oligomerized. Thus, both IDR sequence, and relative degree of oligomerization, contribute to the complex interplay of entropy and enthalpy underlying multiphase condensate organization.

How might the cell have naturally exploited the effects of such differential oligomerization states to evolve biological functions? There is growing evidence that proteins implicated in phase separation feature structured domains that enable binding to RNA or DNA^50,51^. Indeed, RNA binding proteins (RBPs) feature prominently in multiphasic condensates like the nucleolus (NPM1)^20,21,30,52^ and stress granule/P-body (G3BP1/2)^22,53,54^. Truncation mutants of NPM1 lacking either its C-terminal oligomerization domain or N-terminal RNA recognition motif were found to mislocalize when expressed in cells^55^. Similarly, G3BP1 mutants lacking an RNA binding domain (RBDs) were unable to form stress granules when expressed in double KO lines^22^. Furthermore, transcriptional inhibition is well known to result in aberrant nucleolar morphologies and the formation of nucleolar caps^31^. Indeed, it has been suggested that RNA could potentially act as a biological super-scaffold that can tunably drive multiphase formation^56^. Since RBDs can be regulated through post translational modifications, we speculate that living cells could dynamically modulate the oligomerization state of recruited RBDs to drive multiphase condensate formation. Intriguingly, our results also suggest oligomerization as a potential mechanism for altering the spatial localization of condensate components. We speculate that, in the nucleolus, this could potentially be a tunable biophysical mechanism by which fully processed ribosomes are efficiently fluxed out into the nucleoplasm.

The ability to control the miscibility between condensates through the interplay of sequence identity and oligomerization could be exploited for the design of synthetic organelles. Orthogonal scaffolds, potentially actuated externally with light, chemicals, or through other means, could enable the inducible formation of multiphasic condensates for the spatial organization of enzymatic reactions. One possible future strategy, for example, is to explore optogenetic methods that temporally switch miscible components of a single homogenous condensate into demixed components of a spatially segregated, multiphase structure. Such potential engineering applications represent an exciting new frontier, advanced through these and future discoveries, of the increasingly rich ways in which living cells harness oligomerization to spatially organize condensates and control their associated biological functions.

## Methods

### Cell culture and cell-line generation

U2OS (a kind gift from Mark Groudine lab, Fred Hutchinson Cancer Research Center) and Lenti-X 293T (Takara) cells were cultured in a growth medium consisting of Dulbecco’s modified Eagle’s medium (GIBCO), 10% fetal bovine serum (Atlanta Biologicals) and 10□U□ml^−1^ penicillin–streptomycin (GIBCO) and incubated at 37□°C and 5% CO_2_ in a humidified incubator.

### Lentiviral transduction

For Corelet and NPM1 overexpression, lentiviruses containing desired constructs were produced by transfecting the plasmid along with helper plasmids VSVG and PSP (kind gift from Marc Diamond lab, UT Southwestern) into HEK293T cells with Lipofectamine™ 3000 (Invitrogen). Virus was collected 2-3 days after transfection and used to infect WT U2OS. Lentivirus transduction was performed in 96-well plates. Three days following lentivirus application to cells at low confluency, cells were passaged for stable maintenance or directly to 96-well fibronectin-coated glass bottom dishes for live cell microscopy. The infected cells were imaged no earlier than 72□h after infection.

### Yeast plasmid construction

All integration and 2μ plasmids were constructed based on the pJLA vectors using either restriction digest and ligation method with T4 DNA ligase (NEB), or In-Fusion HD cloning kit (Takara Bio). The following restriction enzymes were used for cloning: MreI (Thermo Fisher Scientific), SpeI-HF(NEB), BamHI-HF(NEB), NotI-HF(NEB), AgeI-HF(NEB), AscI (NEB), XhoI (NEB), and SacI-HF(NEB). Promoters (CCW12p, HHF2p, and RPL18Bp) and terminators (TDH1t, ENO2t, and ENO1t) that are not in the pJLA vectors were obtained from a published yeast toolkit on Addgene^49^. Cloned plasmids were transformed into Stellar *E. coli* competent cells (Takara Bio), from which single colonies were isolated from LB plates supplemented with ampicillin. Next, colonies carrying correct clones were identified by colony PCR using OneTaq® Hot Start Quick-Load® 2X Master Mix (NEB), and plasmids were purified from overnight culture in LB containing 150 μg/ml ampicillin at 37°C. All cloned plasmids were verified by Sanger sequencing (Genewiz). Recombinant DNA (I3-01, O3-33, and DDX4_N_CS) were codon-optimized for *S. cerevisiae* using IDT codon optimization tool and purchased as gBlocks® gene fragments from IDT.

### Yeast transformation and culture

Integration plasmids were linearized with PmeI (NEB) and transformed into yeast using the standard lithium acetate method^57^. *S. cerevisiae* CEN.PK2-1C (MATa, ura3-52, trp1-289, leu2-3112, his3Δ1, MAL2-8c, SUC2) was used as the background to construct all yeast strains used in this study. Transformants were selected on synthetic complete (SC) dropout agar plates lacking appropriate amino acid for auxotrophic marker selection, supplemented with 2% (w/v) glucose. To verify integration of linearized DNA construct into the yeast genome, single colonies were isolated and grown in 1mL SC media lacking appropriate amino acid with 2% (w/v) glucose overnight at 30 °C in 24-well plates covered with aluminum foil. In the next day, the overnight culture was diluted 1:50 into fresh media in 24-well plates and grown at 30 °C for 4hr until early exponential phase for imaging (OD_600_ between 1-2). Yeast strains were stored as 20% (v/v) glycerol stocks at -80 °C.

### Airyscan microscopy

The super-resolution fluorescence images were taken with a 100x α Plan-Apochromat 1.46 Oil DIC M27 objective on a Zeiss LSM 980 Airy Scan 2.0 microscope using the Airy Scan SR mode. Imaging was performed using Zeiss Zen Blue 3.2 software. To image multi-channel z-stack image, frame switching was used, and the entire z-stack was imaged per track before switching the channel. The 405nm, 488nm, 561nm, and 639nm lasers were used to image mTagBFP2, GFP, mCherry, and MemBrite® 640/660 Fix dye (Biotium) in cells, respectively. Point Spread Function was verified using TetraSpeck™ microspheres (Thermofisher T7279).

### Quantitative microscopy to determine the valence of NPM1-Corelet

Quantitative microscopy was performed using a Zeiss LSM 980 confocal microscope equipped with a spectral array detector (32-element cooled GaAsP array functioning as a spectral confocal). The spectral array detector enabled fluorescence correlation spectroscopy (FCS) for determining the concentration of fluorophore-tagged proteins (GFP and mCherry) in live cells. First, the effective confocal volume (*V*_eff_) of the objective (60 X, 1.43 N. A., oil, structural parameter = 7) was determined at a z-height close to the common cell imaging plane. Following the established protocol^58^, the *V*_eff_ when using 488 nm and 561 nm lasers were determined with aqueous solutions of Atto 488 (concentration = 200 nM) and Alexa Fluor 568 (concentration = 50 nM), respectively at 37 °C.

In cells that stably express GFP-P2A-mCherry, we used FCS to determine the number of diffusing particles in a corresponding *V*_eff_. Using the *V*_eff_ measured using dye solutions, the concentration of GFP and mCherry were determined. A series of images were then collected using Zeiss LSM online fingerprinting with a wide range of laser power (0.01-5%) and gain (650-850 V). Using the image intensity at different acquisition settings, intensity-concentration calibration curves were determined (at least 3 different cells for each fluorophore). The concentration of NPM1-mCh-SspB and NLS-iLID-GFP-FTH1 were determined from these calibration curves, similar to previously reported methods^39^. All the multicolor mammalian cell images and time series were collected using the spectral unmixing of mTagBFP2, GFP, and mCherry using the online fingerprinting module on the Zeiss LSM. The optogenetic activation was performed with lasers 405 nm (intensity = 0.2%) and 488 nm (intensity = 0.3%).

### Yeast sample preparation for fixed-cell imaging

For fixed-cell imaging experiments on Zeiss LSM 980 with Airyscan, strains were streaked from glycerol stocks, cultured as described above, and imaged on the 96-well plate. The wells were coated with 50 μl solution of 1 mg/ml concanavalin A (ConA) (Sigma-Aldrich L7647) in 20 mM NaOAc for yeast immobilization, as previously described^59^. The yeast culture was diluted to an OD_600_ of 1, and 100 μl of the diluted culture was added to each well. Yeast surface staining using the MemBrite® 640/660 Fix dye was done after immobilization and prior to fixation following the manufacture protocol. Yeast was fixed with 100 μl 4% PFA in PBS at room temperature for 20 min. Finally, each well was washed twice with PBS and 100 μl PBS was added before imaging.

### Normalized center-to-center distance calculation

The condensate sizes and distances in yeast imaging experiments were measured using the Distance Analysis (DiAna) plugin in Fiji ImageJ2^60,61^. The condensates in each channel were segmented using the iterative threshold method with a minimum size of 50 pixels and a minimum threshold of 20% of the average maximum intensity across images of the same well. Condensate sizes and the center-to-center distances between overlapping condensates were obtained using the DiAna colocalization analysis. Outputs of the DiAna colocalization analysis were individually verified. The center-to-center distance between overlapping condensates was then normalized by the sum of radii of the two condensates. To account for noise in the measurements, the bin width was set to 0.1 in the frequency distribution.

### Simulation Methods

Direct coexistence simulations were performed to estimate the relative miscibility of SV polyampholyte pairs^35,37^. To model molecular interactions between the monomers, we utilized the HPS-implicit solvent model^62^. Multivalency in polyampholytes was modelled in our simulations by covalently linking polyampholytes (using HPS bond parameters) to a central hard-sphere bead with diameter 1.2 Å, roughly double the diameter of a lysine or glutamate bead. All simulations runs were performed using HOOMD-Blue^63^.

Initial configurations were generated by randomly placing both polyampholyte species, at equal mass fraction, in a 50×50×50 nm cubic box and performing a short run in which the simulation box was compressed down to either 20×20×20 nm or 25×25×25 nm, depending on system size and valence of polyampholytes. The system sizes used for different valence states are described in supplementary table S1. In accordance with the direct coexistence method^64,65^, the z-dimension of the box was then expanded to 125 nm by adding empty space to either side, and an equilibration run was performed at constant NVT. Finally, using the equilibrated NVT configurations, NPAT (keeping pressure, temperature and cross-sectional area of box constant) direct coexistence runs were performed to obtain the miscibility behavior. A timestep of 10 fs was used for all runs and both the NVT and NPAT simulations were run for 5 μs. All simulations were run at a temperature of 250 K which is lower than the critical temperature of sequence SV1 (the polyampholyte sequence with the lowest critical temperature). For NVT simulations, a Langevin thermostat with a friction coefficient of 1ps^-1^ was used while for NPAT simulations, an MTK thermostat-barostat was used with coupling constants *τ* = 0.5 and *τ*_*p*_ 0.5.

To obtain coexistence data, density profiles for both polyampholyte species were recorded along the z-dimension and averaged over the entire trajectory. For estimating the partitioning, we fit a hyperbolic tangent function to the density profile to estimate the concentration of the dense and dilute phases, as well as the spatial extent of each phase in the z-dimension. Partitioning was then estimated as described previously. Interfacial tension (Fig 6B) was calculated from the components of the pressure tensor as previously described in the literature^65,66^. All simulation snapshots were generated using the “freud” python library^67^.

## Acknowledgements

We thank Anita Donlic and Grace Bechtel for early helpful discussions, David Sanders for the FM5 vectors, Evangelos Gatzogiannis for microscopy assistance, and other members of the Brangwynne Lab for helpful feedback and discussions.

This work was supported by the Howard Hughes Medical Institute, the Princeton Biomolecular Condensate Program, the AFOSR MURI (FA9550-20-1-0241), the U.S. Department of Energy, Office of Science, Office of Biological and Environmental Research (DE-SC0022155), and the Princeton Center for Complex Materials (PCCM), a U.S. National Science Foundation Materials Research Science and Engineering Center (Grant Nos. DMR-1420541 and DMR-2011750). Simulations were performed using computational resources provided by the Princeton Institute for Computational Science and Engineering (PICSciE) and the Office of Information Technology’s High Performance Computing Center and Visualization Laboratory at Princeton University. A.N. is a Howard Hughes Medical Institute Awardee of the Life Sciences Research Foundation. M.T.W. is supported by the NSF GRFP (DGE-1656466).

## Contributions

U.R., K.X., A.Z.P, J.L.A and C.P.B. designed the study. U.R., K.X., A.N., M.T.W. performed research and contributed reagents. U.R. and K.X. analyzed the data. U.R., K.X., A.Z.P., J.L.A and C.P.B. wrote the manuscript with contributions from all authors.

