## SupplementaryInformation for "Asymmetric oligomerization state and sequence patterning can tune multiphase condensate miscibility"

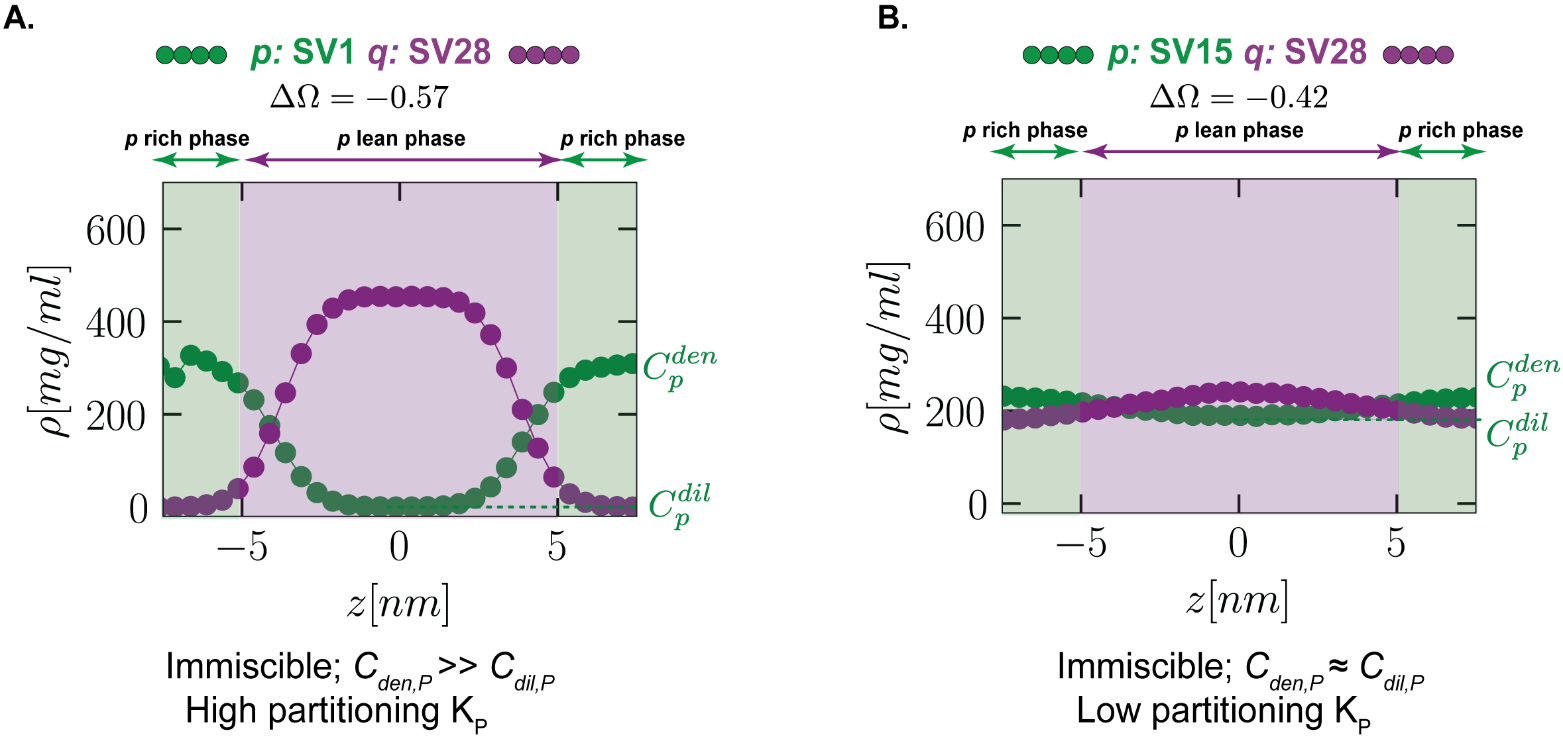

**Supplementary Fig. 1. Partitioning metric as a proxy for immiscibility.** **a.** Density profiles of immiscible SV1 and SV28 showing the extent of the two phases and the high partitioning. **b.** Density profiles of relatively miscible SV15 and SV28 showing marginal demixing between the two phases and the low partitioning.

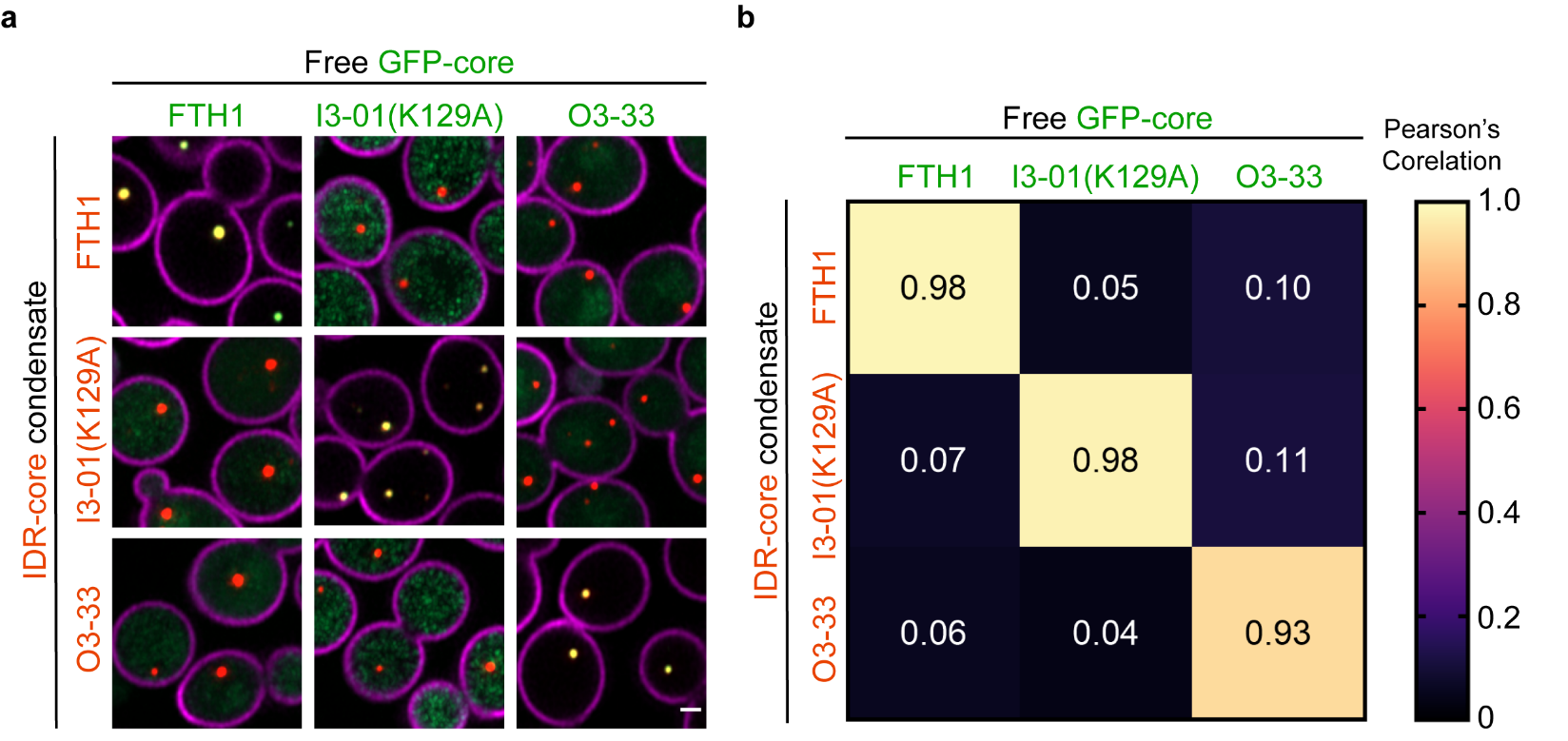

**Supplementary Fig. 2. Additional data demonstrate core orthogonality Related to Fig. 2.** **a.** Confocal fluorescence microscope images of yeast expressing combinations of mCherry-tagged IDR-core condensate and GFP-tagged free core. Scale bars, 0.5 μm. **b.** The degree of recruitment of free core into the condensate as measured using Pearson’s correlation.

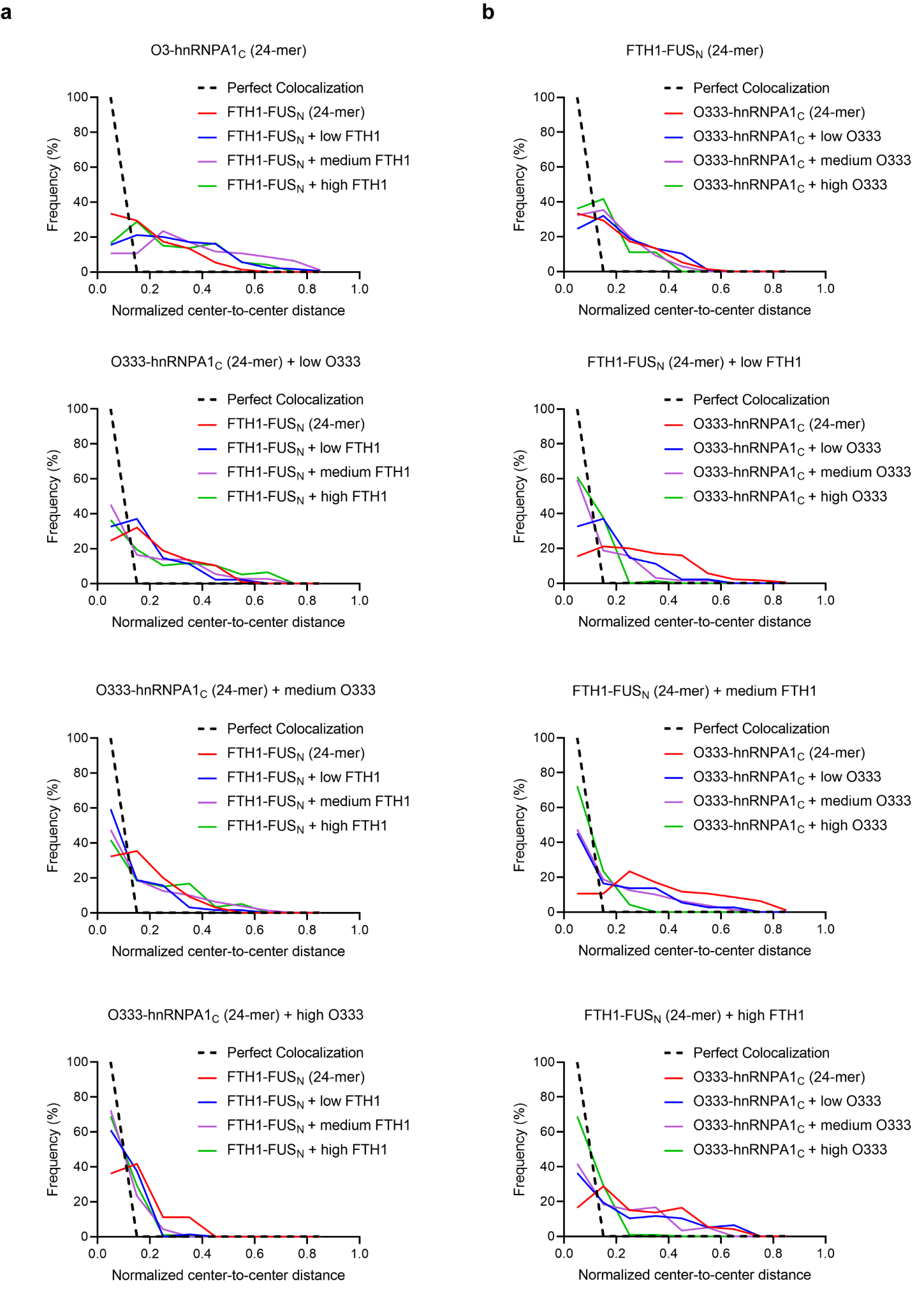

**Supplementary Fig. 3. The normalized center-to-center distribution between FUS_N_-FTH1 and hnRNPA1_C_-O3-33 condensates Related to Fig. 3. a.** Varying FUS_N_ valence at a fixed hnRNPA1_C_ valence. **b.** Varying hnRNPA1_C_ valence at a fixed FUS_N_ valence.

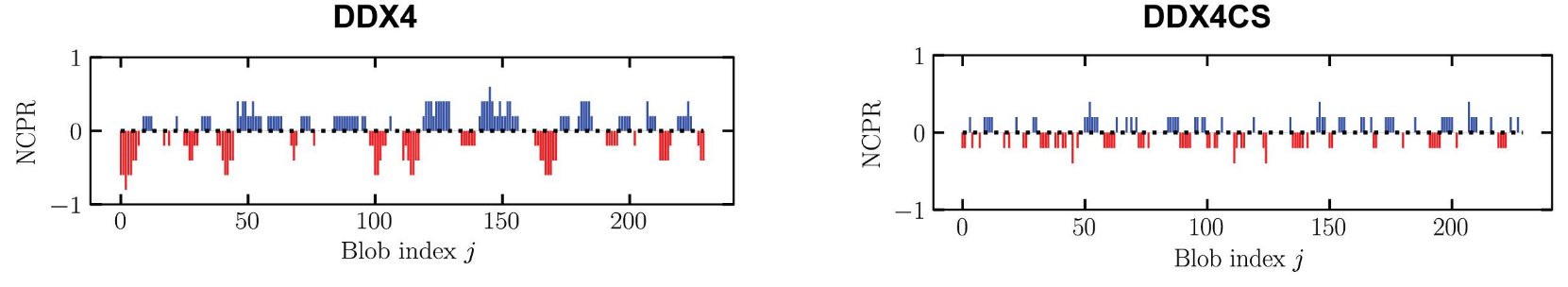

**Supplementary Fig. 4. Comparison of the net charge per residue (NCPR) between DDX4_N_ and DDX4_N_CS.** The charges amino acid blocks in DDX4_N_ are scrambled to obtain DDX4_N_CS.

**Supplementary Table 1. System sizes for simulations**

| Valence | No. of oligomerized polyampholytes *p* molecules | No. of un-oligomerized  polyampholytes *q* molecules |
| --- | --- | --- |
| 1 | 375 | 375 |
| 3 | 147 | 441 |
| 5 | 108 | 529 |
| 7 | 108 | 729 |

**Supplementary Table 2. Yeast strains used in this study.**

| Strain | Description | Genotype | Source |
| --- | --- | --- | --- |
| CEN.PK2-1C | Wild-type *Saccharomyces cerevisiae* | *MATa ura3-52 trp1-289 leu2-3,112 his3-1 MAL2-8c SUC2* | ^1^ |
| yKX371 | Investigate core orthogonality | *his3::HIS3*-TEF1p_FUS_N_-mCherry-FTH1_TDH1t  *leu2::LEU2*-PGK1p_GFP-FTH1_CYC1t | This study |
| yKX372 | Investigate core orthogonality | *his3::HIS3*-TEF1p_FUS_N_-mCherry-FTH1_TDH1t  *leu2::LEU2*-PGK1p_GFP-I301(K129A)_ENO2t | This study |
| yKX373 | Investigate core orthogonality | *his3::HIS3*-TEF1p_FUS_N_-mCherry-FTH1_TDH1t *leu2::LEU2*-PGK1p_O333-GFP_ENO2t | This study |
| yKX374 | Investigate core orthogonality | *his3::HIS3*-TEF1p_FUS_N_-mCherry-I301(K129A)_TDH1t *leu2::LEU2*-PGK1p_GFP-FTH1_CYC1t | This study |
| yKX375 | Investigate core orthogonality | *his3::HIS3*-TEF1p_FUS_N_-mCherry-I301(K129A)_TDH1t *leu2::LEU2*-PGK1p_GFP-I301(K129A)_ENO2t | This study |
| yKX376 | Investigate core orthogonality | *his3::HIS3*-TEF1p_FUS_N_-mCherry-I301(K129A)_TDH1t *leu2::LEU2*-PGK1p_O333-GFP_ENO2t | This study |
| yKX377 | Investigate core orthogonality | *his3::HIS3*-TEF1p_O333-mCherry-hnRNPA1_C__TDH1t *leu2::LEU2*-PGK1p_GFP-FTH1_CYC1t | This study |
| yKX378 | Investigate core orthogonality | *his3::HIS3*-TEF1p_O333-mCherry-hnRNPA1_C__TDH1t *leu2::LEU2*-PGK1p_GFP-I301(K129A)_ENO2t | This study |
| yKX379 | Investigate core orthogonality | *his3::HIS3*-TEF1p_O333-mCherry-hnRNPA1_C__TDH1t *leu2::LEU2*-PGK1p_O333-GFP_ENO2t | This study |
| yKX172 | Synthetic IDR-core proteins immiscibility | *his3::HIS3*-TEF1p_FUS_N_-mCherry-I301(K129A)_TDH1t  *leu2::LEU2*-PGK1p_FUS_N_-GFP-FTH1_ENO2t | This study |
| yKX173 | Synthetic IDR-core proteins immiscibility | *his3::HIS3*-TEF1p_DDX4­_N_-mCherry-I301(K129A)_TDH1t *leu2::LEU2*-PGK1p_FUS_N_-GFP-FTH1_ENO2t | This study |
| yKX294 | Synthetic IDR-core proteins immiscibility | *his3::HIS3*-TEF1p_DDX4­_N_CS-mCherry-I301(K129A)_TDH1t *leu2::LEU2*-PGK1p_FUS_N_-GFP-FTH1_ENO2t | This study |
| yKX175 | Synthetic IDR-core proteins immiscibility | *his3::HIS3*-TEF1p_FUS_N_-mCherry-I301(K129A)_TDH1t *leu2::LEU2*-PGK1p_DDX4_N_-GFP-FTH1_ENO2t | This study |
| yKX176 | Synthetic IDR-core proteins immiscibility | *his3::HIS3*-TEF1p_DDX4_N_-mCherry-I301(K129A)_TDH1t  *leu2::LEU2*-PGK1p_DDX4_N_-GFP-FTH1_ENO2t | This study |
| yKX297 | Synthetic IDR-core proteins immiscibility | *his3::HIS3*-TEF1p_DDX4_N_CS-mCherry-I301(K129A)_TDH1t *leu2::LEU2*-PGK1p_DDX4_N_-GFP-FTH1_ENO2t | This study |
| yKX296 | Synthetic IDR-core proteins immiscibility | *his3::HIS3*-TEF1p_FUS_N_-mCherry-I301(K129A)_TDH1t  *leu2::LEU2*-PGK1p_DDX4_N_CS-GFP-FTH1_ENO2t | This study |
| yKX331 | Synthetic IDR-core proteins immiscibility | *his3::HIS3*-TEF1p_DDX4_N_CS-mCherry-I301(K129A)_TDH1t *leu2::LEU2*-PGK1p_DDX4_N_CS-GFP-FTH1_ENO2t | This study |
| yKX332 | Synthetic IDR-core proteins immiscibility | *his3::HIS3*-TEF1p_DDX4_N_-mCherry-I301(K129A)_TDH1t *leu2::LEU2*-PGK1p_DDX4_N_CS-GFP-FTH1_ENO2t | This study |
| Strain | Description | Genotype | Source |
| yKX335 | Oligomerization-driven immiscibility | *his3::HIS3*-TEF1p_O333-mCherry-hnRNPA1_C__TDH1t *leu2::LEU2*-PGK1p_FUS_N_-GFP-FTH1_ENO2t_ Rev(TEF1p_mTagBFP2-FTH1_ACT1t) | This study |
| yKX336 | Oligomerization-driven immiscibility | *his3::HIS3*-TEF1p_O333-mCherry-hnRNPA1_C__TDH1t *leu2::LEU2*-PGK1p_FUS_N_-GFP-FTH1_ENO2t_ Rev(TEF1p_mTagBFP2-FTH1_ACT1t) *trp1::TRP1*-CCW12p_mTagBFP2-FTH1_ENO1t | This study |
| yKX337 | Oligomerization-driven immiscibility | *his3::HIS3*-TEF1p_O333-mCherry-hnRNPA1_C__TDH1t *leu2::LEU2-*PGK1p_FUS_N_-GFP-FTH1_ENO2t_ Rev(TEF1p_mTagBFP2-FTH1_ACT1t) *trp1::TRP1*-CCW12p_mTagBFP2-FTH1_ENO1t_ Rev(TEF1p_mTagBFP2-FTH1_ACT1t) | This study |
| yKX349 | Oligomerization-driven immiscibility | *his3::HIS3*-TEF1p_O333-mCherry-hnRNPA1_C__TDH1t_ Rev(TDH3p_O333-mCherry_ENO1t) *leu2::LEU2*-PGK1p_FUS_N_-GFP-FTH1_ENO2t_ Rev(TEF1p_mTagBFP2-FTH1_ACT1t) | This study |
| yKX355 | Oligomerization-driven immiscibility | *his3::HIS3*-TEF1p_O333-mCherry-hnRNPA1_C__TDH1t_ Rev(TDH3p_O333-mCherry_ENO1t) *leu2::LEU2*-PGK1p_FUS_N_-GFP-FTH1_ENO2t_ Rev(TEF1p_mTagBFP2-FTH1_ACT1t) *trp1::TRP1*-CCW12p_mTagBFP2-FTH1_ENO1t | This study |
| yKX356 | Oligomerization-driven immiscibility | *his3::HIS3*-TEF1p_O333-mCherry-hnRNPA1_C__TDH1t_ Rev(TDH3p_O333-mCherry_ENO1t) *leu2::LEU2*-PGK1p_FUS_N_-GFP-FTH1_ENO2t_ Rev(TEF1p_mTagBFP2-FTH1_ACT1t) *trp1::TRP1*-CCW12p_mTagBFP2-FTH1_ENO1t_ Rev(TEF1p_mTagBFP2-FTH1_ACT1t) | This study |
| yKX358 | Oligomerization-driven immiscibility | *his3::HIS3*-TEF1p_O333-mCherry-hnRNPA1_C__TDH1t_ Rev(HHF2p_O333-mCherry_ENO1t) *leu2::LEU2*-PGK1p_FUS_N_-GFP-FTH1_ENO2t_ Rev(TEF1p_mTagBFP2-FTH1_ACT1t) | This study |
| yKX359 | Oligomerization-driven immiscibility | *his3::HIS3*-TEF1p_O333-mCherry-hnRNPA1_C__TDH1t_ Rev(HHF2p_O333-mCherry_ENO1t) *leu2::LEU2*-PGK1p_FUS_N_-GFP-FTH1_ENO2t_ Rev(TEF1p_mTagBFP2-FTH1_ACT1t) *trp1::TRP1*-CCW12p_mTagBFP2-FTH1_ENO1t | This study |
| yKX360 | Oligomerization-driven immiscibility | *his3::HIS3*-TEF1p_O333-mCherry-hnRNPA1_C__TDH1t_ Rev(HHF2p_O333-mCherry_ENO1t) *leu2::LEU2*-PGK1p_FUS_N_-GFP-FTH1_ENO2t_ Rev(TEF1p_mTagBFP2-FTH1_ACT1t) *trp1::TRP1*-CCW12p_mTagBFP2-FTH1_ENO1t_ Rev(TEF1p_mTagBFP2-FTH1_ACT1t) | This study |
| yKX362 | Oligomerization-driven immiscibility | *his3::HIS3*-TEF1p_O333-mCherry-hnRNPA1c_TDH1t_ Rev(RPL18Bp_O333-mCherry_ENO1t) *leu2::LEU2*-PGK1p_FUSn-GFP-FTH1_ENO2t_ Rev(TEF1p_mTagBFP2-FTH1_ACT1t) | This study |
| Strain | Description | Genotype | Source |
| yKX363 | Oligomerization-driven immiscibility | *his3::HIS3*-TEF1p_O333-mCherry-hnRNPA1c_TDH1t_ Rev(RPL18Bp_O333-mCherry_ENO1t) *leu2::LEU2*-PGK1p_FUSn-GFP-FTH1_ENO2t_ Rev(TEF1p_mTagBFP2-FTH1_ACT1t) *trp1::TRP1*-CCW12p_mTagBFP2-FTH1_ENO1t | This study |
| yKX364 | Oligomerization-driven immiscibility | *his3::HIS3*-TEF1p_O333-mCherry-hnRNPA1c_TDH1t_ Rev(RPL18Bp_O333-mCherry_ENO1t) *leu2::LEU2*-PGK1p_FUSn-GFP-FTH1_ENO2t_ Rev(TEF1p_mTagBFP2-FTH1_ACT1t) *trp1::TRP1*-CCW12p_mTagBFP2-FTH1_ENO1t_ Rev(TEF1p_mTagBFP2-FTH1_ACT1t) | This study |

**Supplementary Table 3. Plasmids used in this study.**

| Plasmid | Description | Source |
| --- | --- | --- |
| **Yeast** | | |
| pKX020 | Amp^R^, HIS3 integration, TEF1p_DDX4_N_CS-mCherry-I301(K129A)_TDH1t | This study |
| pKX100 | Amp^R^, HIS3 integration, TEF1p_FUSn-mCherry-FTH1_TDH1t | This study |
| pKX101 | Amp^R^, LEU2 integration, PGK1p_GFP-FTH1_CYC1t | This study |
| pKX117 | Amp^R^, HIS3 integration, TEF1p_FUS_N_-mCherry-I301(K129A)_TDH1t | This study |
| pKX119 | Amp^R^, HIS3 integration, TEF1p_DDX4_N_-mCherry-I301(K129A)_TDH1t | This study |
| pKX121 | Amp^R^, LEU2 integration, PGK1p_FUS_N_-GFP-FTH1_ENO2t | This study |
| pKX123 | Amp^R^, LEU2 integration, PGK1p_DDX4_N_-GFP-FTH1_ENO2t | This study |
| pKX180 | Amp^R^, LEU2 integration, PGK1p_GFP-I301(K129A)_ENO2t | This study |
| pKX328 | Amp^R^, HIS3 integration, TEF1p_O333-mCherry-hnRNPA1_C__TDH1t | This study |
| pKX338 | Amp^R^, LEU2 integration, PGK1p_DDX4_N_CS-GFP-FTH1_ENO2t | This study |
| pKX393 | Amp^R^, 2μ, URA3, TEF1p_mTagBFP2-FTH1_ACT1t | This study |
| pKX394 | Amp^R^, LEU2 integration, PGK1p_FUS_N_-GFP-FTH1_ENO2t_ Rev(TEF1p_mTagBFP2-FTH1_ACT1t) | This study |
| pKX395 | Amp^R^, TRP1 integration, CCW12p_mTagBFP2-FTH1_ENO1t | This study |
| pKX396 | Amp^R^, TRP1 integration, CCW12p_mTagBFP2-FTH1_ENO1t_ Rev(TEF1p_mTagBFP2-FTH1_ACT1t) | This study |
| pKX399 | Amp^R^, HIS3 integration, TDH3p_O333-mCherry_ENO1t | This study |
| pKX400 | Amp^R^, HIS3 integration, TEF1p_O333-mCherry-hnRNPA1_C__TDH1t_ Rev(TDH3p_O333-mCherry_ENO1t) | This study |
| pKX430 | Amp^R^, HIS3 integration, HHF2p_O333-mCherry_ENO1t | This study |
| pKX431 | Amp^R^, HIS3 integration, RPL18Bp_O333-mCherry_ENO1t | This study |
| pKX433 | Amp^R^, HIS3 integration, TEF1p_O333-mCherry-hnRNPA1_C__TDH1t_ Rev(HHF2p_O333-mCherry_ENO1t) | This study |
| pKX434 | Amp^R^, HIS3 integration, TEF1p_O333-mCherry-hnRNPA1_C__TDH1t_ Rev(RPL18Bp_O333-mCherry_ENO1t) | This study |
| pKX452 | Amp^R^, LEU2 integration, PGK1p_O333-GFP_ENO2t | This study |
| **Mammalian** | | |
|  | FM5::NPM1-mCherry-sspB | This study |
|  | FM5::NLS-iLID-GFP-FTH1 | This study |
|  | FM5::mTagBFP2-NPM1 | This study |
